# The Stone Age Plague: 1000 years of Persistence in Eurasia

**DOI:** 10.1101/094243

**Authors:** Aida Andrades Valtueña, Alissa Mittnik, Felix M. Key, Wolfgang Haak, Raili Allmäe, Andrej Belinskij, Mantas Daubaras, Michal Feldman, Rimantas Jankauskas, Ivor Janković, Ken Massy, Mario Novak, Saskia Pfrengle, Sabine Reinhold, Mario Šlaus, Maria A. Spyrou, Anna Szecsenyi-Nagy, Mari Tõrv, Svend Hansen, Kirsten I. Bos, Philipp W. Stockhammer, Alexander Herbig, Johannes Krause

## Abstract

Molecular signatures of *Yersinia pestis* were recently identified in prehistoric Eurasian individuals, thus suggesting *Y. pestis* caused some form of disease in humans prior to the first historically documented pandemic. Here, we present six new *Y. pestis* genomes spanning from the European Late Neolithic to the Bronze Age (LNBA) dating from 4,800 to 3,700 BP. We show that all currently investigated LNBA strains form a single genetic clade in the *Y. pestis* phylogeny that appears to be extinct. Interpreting our data within the context of recent ancient human genomic evidence, which suggests an increase in human mobility during the LNBA, we propose a possible scenario for the spread of *Y. pestis* during the LNBA: *Y. pestis* may have entered Europe from Central Eurasia during an expansion of steppe people, persisted within Europe until the mid Bronze Age, and moved back towards Central Eurasia in parallel with subsequent human population movements.

## Introduction

Plague pandemics throughout human history caused unprecedented levels of mortality that contributed to profound socioeconomic and political changes. Conventionally it is assumed that plague affected human populations in three pandemic waves. The first, the Plague of Justinian, starting in the 6th century AD, was followed by multiple epidemic outbreaks in Europe and the Mediterranean basin, and has been associated with the weakening and decay of the Byzantine empire (Russell, 1968). The second plague pandemic first struck in the 14th century with the infamous ‘Black Death’ (1347-1352), which again spread from Asia to Europe seemingly along both land and maritime routes (Zietz and Dunkelberg, 2004). It is estimated that this initial onslaught killed 50% of the European population (Benedictow, 2004). It was followed by outbreaks of varying intensity that lasted until the late eighteenth century (Cohn JR, 2008). The most recent plague pandemic started in the 19th century and began in the Yunnan province of China. It reached Hong Kong by 1894 and followed global trade routes to achieve a near worldwide distribution (Stenseth et al., 2008). Since then plague has persisted in rodent populations in many areas of the world and continues to cause both isolated human cases and local epidemics (http://www.who.int/mediacentre/factsheets/fs267/en/).

Plague is caused by a systemic infection with the Gram-negative bacterium *Yersinia pestis.* Advances in ancient DNA (aDNA) research have permitted the successful reconstruction of a series of *Y. pestis* genomes from victims of both the first and second plague pandemic, thus confirming a *Y. pestis* involvement and providing new perspectives on how this bacterium historically spread through Europe (Bos et al., 2016, 2011; Feldman et al., 2016; Spyrou et al., 2016; Wagner et al., 2014). Most recently, a study by Spyrou et al. (2016) suggested that during the second pandemic, a European focus was established from where subsequent outbreaks, such as the Ellwangen outbreak (16^th^ century Germany) or the Great Plague of Marseille (1720-1722 France), were derived (Spyrou et al., 2016). The authors also proposed that a descendant of the Black Death strain travelled eastwards in the late 14^th^ century, became established in East Asia and subsequently gave rise to the most recent plague pandemic that spread the pathogen around the globe.

Our perception of the evolutionary history of *Y. pestis* was changed substantially by a recent report of two reconstructed genomes from Bronze Age individuals found in the Altai region (Southern Siberia, dating to ∼4,729 cal BP and ∼3,635 cal BP, respectively) and molecular *Y. pestis* signatures in an additional five individuals from Eurasia (∼4,500 to 2,800 BP) suggesting the presence of plague in human populations over a diffuse geographic range prior to the first historically recorded pandemics. Phylogenetic analysis of the two reconstructed *Y. pestis* genomes from the Altai region shows that they occupy a phylogenetic position ancestral to all extant *Y. pestis* strains, though this branch was not adequately resolved (bootstrap lower than 95% (Rasmussen et al., 2015)). Further open questions remain regarding *Y. pestis*' early association with humans. It is not currently known whether the *Y. pestis* lineages circulating in Europe during the Late Neolithic and Bronze Age were all descended from the ∼5,000 BP Central Eurasian strain or whether there were multiple strains circulating in Europe and Asia. Furthermore, how did plague spread over such a vast territory during the period comprising the Late Neolithic and Bronze Age? Could these bacterial strains have been associated with certain human groups and their respective subsistence strategies and cultures?

The Late Neolithic and Early Bronze Age in Western Eurasia (ca. 4,900-3,700/3,600 BP; cf. Stockhammer et al., 2015) was a time of major transformative cultural and social changes that led to cross-European networks of contact and exchange (Vandkilde, 2016). Intriguingly, recent studies on ancient human genomes suggested a major expansion of people from the Eurasian Steppe westwards into Central Europe as well as eastwards into Central Eurasia and Southern Siberia starting around 4,800 BP (Allentoft et al., 2015; Haak et al., 2015). These steppe people with a predominantly pastoral economy carried a genetic component that is present in all Europeans today but was absent in early and middle Neolithic farmers in Europe prior to their arrival. The highest amount of genetic ‘steppe ancestry’ in ancient Europeans was found in individuals associated with the Late Neolithic Corded Ware Complex (Figure 1) around 4,500 BP (Haak et al., 2015), who show a genetic makeup close to the steppe people associated with the ‘Yamnaya’ complex, suggesting a strong genetic link between those two groups. Furthermore, it could be shown that the ‘middle Neolithic farmer’ genetic component also appears in individuals associated with the Andronovo culture in the Altai region around 4,200 BP (Allentoft et al., 2015). These genetic links between humans ranging from Western Eurasia to Southern Siberia highlight the dimensions of mobility and connectedness at the time of the Bronze Age.

**Figure 1:**
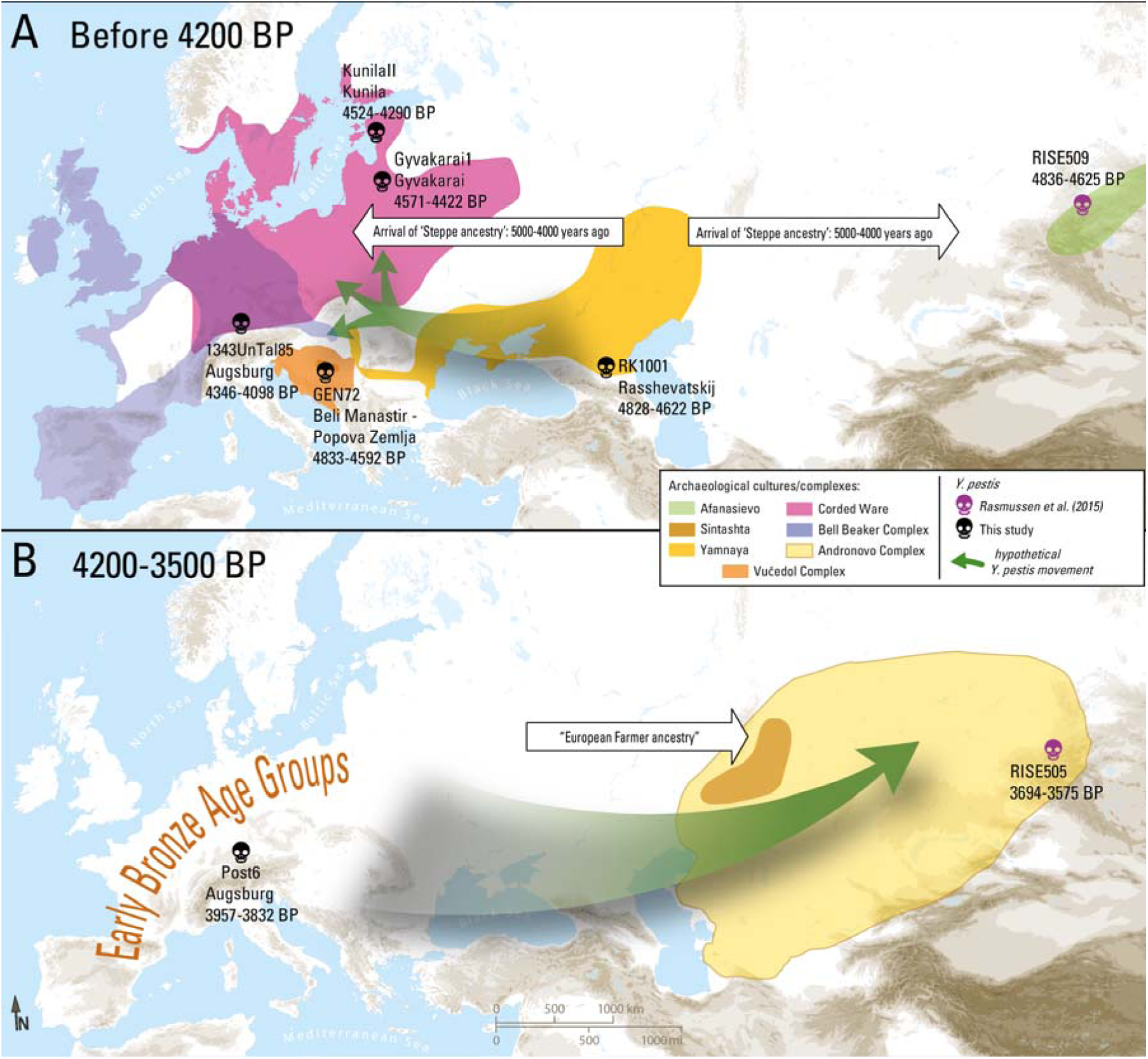
Map of proposed Yersinia pestis circulation throughout Eurasia. A) Entrance of Y. pestis into Europe from Central Eurasia with the expansion of Yamnaya pastoralists around 4,800 years ago. B) Circulation of Y.pestis back into the Altai from Europe. Only complete genomes are shown.

The reasons for the magnitude of the genetic turnover that occurred in Central Europe around 4800 BP, where around 75% of the local middle Neolithic farmer genetics was replaced (Haak et al., 2015), have yet to be explained. As in other episodes of human history, infectious diseases may have played a significant role in triggering or catalyzing those major cultural shifts and human migrations. Here we present six novel *Y. pestis* genomes from Central Europe and the North Caucasus steppe spanning from the Late Neolithic to the Bronze Age (LNBA). Through comparative analyses with other ancient and modern *Y. pestis* lineages (Bos et al., 2016, 2011; Cui et al., 2013; Feldman et al., 2016; Kislichkina et al., 2015; Spyrou et al., 2016; Zhgenti et al., 2015), we show that all LNBA strains form a single clade in the *Y. pestis* phylogeny. This indicates a common origin of all currently identified *Y. pestis* strains circulating in Eurasia during the Late Neolithic and Bronze Age, and reveals a distribution pattern that parallels human movements in time and space.

## Results

### Screening

A total of 563 tooth and bone samples dating from the Late Neolithic to the Bronze Age from Russia (122), Hungary and Croatia (139), Lithuania (27), Estonia (45), Latvia (10), and Germany (Althausen 4, Augsburg 83, Mittelelbe-Saale 133) were screened for *Y. pestis* by mapping DNA sequencing reads ranging in numbers from 700,000 to 21,000,000 against a multi-fasta reference consisting of the genomes of 12 different *Yersinia* species (Table 1).

**Table 1:** Genomes from the NCBI (RefSeq/Nucleotide) database, used in the multi-species reference panel for screening for *Y. pestis* aDNA.

| Species name | Strain | NCBI Accession number |
| --- | --- | --- |
| *Y. pestis* | CO92 | NC_003143.1 |
| *Y. pseudotuberculosis* | IP 32953 | NC_006155.1 |
| *Y. enterocolitica* | subsp. enterocolitica 8081 | NC_008800.1 |
| *Y. aldovae* | ATCC 35236 | NZ_ACCB01000210.1 |
| *Y. bercovieri* | ATCC 43970 | NZ_AALC02000229.1 |
| *Y. frederiksenii* | ATCC 33641 | NZ_AALE02000161.1 |
| *Y. intermedia* | ATCC 29909 | NZ_AALF02000123.1 |
| *Y. kristensenii* | ATCC 33638 | NZ_ACCA01000153.1 |
| *Y. mollaretii* | ATCC 43969 | NZ_AALD02000179.1 |
| *Y. rohdei* | ATCC 43380 | NZ_ACCD01000141.1 |
| *Y. ruckeri* | ATCC 29473 | NZ_ACCC01000174.1 |

To assess if an individual was positive for *Y. pestis*, we calculated a score based on the number of specific reads mapping to *Y. pestis* in comparison to the number of reads mapping to other *Yersinia* species (See methods). Following this metric, all individuals with a positive score were identified as possible candidates. Individuals that had a score higher than 0.005, and had reads mapping to all the three plasmids present in *Y. pestis* were considered ‘strong’ positives. In our dataset we identified five strong candidates, all of them tooth samples, from three different locations spanning from the Late Neolithic to the Early Bronze Age: one individual from the site Rasshevatskiy (RK1001) in the North Caucasus (Russia), one individual from the Lithuanian site Gyvakarai (Gyvakarai1), one individual from the Estonian site Kunila (KunilaII) and two individuals from Augsburg, Germany (Haunstetten, Unterer Talweg 85 Feature 1343 (1343UnTal85), and Haunstetten, Postillionstraße Feature 6 (6Post). Additionally, one individual from the Croatian site Beli Manastir - Popova Zemlja (GEN72, also a tooth sample), which did not pass the criteria for a strong candidate, was taken along as a potential candidate since it had the highest number of reads mapping to the *Y. pestis* chromosome and all the plasmids (chromosome=993, pCD1=243, pMT1=111, pPCP1=22). For a detailed description of all samples, individuals and archaeological sites see Table 2 and SI.

**Table 2:** Statistic of the *Y. pestis* genome reconstruction. BP = Before Present.

| Individual | Tissue sampled | Site | Country | Dating (Median cal BP) | In-solution Capture | Clipped, merged and quality-filtered reads before mapping | Unique reads mapping to *Y. pestis* reference | Endogenous DNA (%) | Mean Coverage | Coverage (%) |  |  |  |  |
| --- | --- | --- | --- | --- | --- | --- | --- | --- | --- | --- | --- | --- | --- | --- |
|  |  |  |  |  |  |  |  |  |  | >=1X | >=2X | >=3X | >=4X | >=5X |
| RK1001 | Tooth | Rasshevatskiy | Russia | 4720 | no | 1,529,935,532 | 119,540 | 0.01 | 1.0213 | 58.11 | 27.24 | 10.87 | 3.90 | 1.32 |
|  |  |  |  |  | yes | 303,148,884 | 383,900 | 0.85 | 3.3984 | 82.83 | 69.42 | 55.3 | 41.95 | 30.36 |
|  |  |  |  |  | Combined shotgun/capture | 1,833,084,416 | 418,581 | 0.17 | 3.6816 | 86.16 | 73.67 | 59.63 | 45.88 | 33.76 |
| GEN72 | Tooth | Beli Manastir-Popova Zemlja | Croatia | 4721 | yes | 19,777,683 | 1,321,320 | 24.36 | 12.6549 | 91.65 | 89.15 | 86.61 | 83.84 | 80.85 |
| Gyvakarai1 | Tooth | Gyvakarai | Lithuania | 4427 | no | 1,021,452,137 | 473,207 | 0.05 | 5.2245 | 94.07 | 90.96 | 84.12 | 73.1 | 59.1 |
| Kunilall | Tooth | Kunila | Estonia | 4203 | no | 379,155,741 | 546,243 | 0.16 | 5.5418 | 92.48 | 86.65 | 77.58 | 66.49 | 54.77 |
| 1343UnTal85 | Tooth | Augsburg | Germany | 3873 | no | 1,174,989,269 | 1,165,435 | 0.14 | 10.5745 | 93.69 | 93.29 | 92.59 | 91.29 | 89,.04 |
| 6Post | ooth | ugsburg | Germany | 3635 | o | 419,717,299 | 598,030 | 0.17 | 5.3062 | 89.71 | 81.66 | 71.14 | 59.82 | 48.99 |

### Genome reconstruction

The five strong positive individuals identified during the screening step (Gyvakarai1, KunilaII, 1343Untal85, 6Post, RK1001) were shotgun sequenced to a depth of 379,155,741 to 1,529,935,532 reads. In addition to the shotgun sequencing, RK1001 and GEN72 were enriched for *Y. pestis* DNA following an in-solution approach (See Methods). After mapping to the reference genome (*Y. pestis* CO92, NC_003143.1), we reconstructed genomes from all six potential candidates with a mean coverage from 3.7 to 12-fold with 86-94% of the reference genome covered at least 1-fold (Table 2). The reads were independently mapped to the three plasmids of *Y. pestis* CO92, and we reconstructed the three plasmids for our ancient samples with mean coverage of: pCD1 7 to 24-fold, pMT1 3 to 14-fold and pPCP1 18 to 43-fold (Supplementary Table 1).

In order to authenticate the ancient origin of the bacterial genomes, we evaluated the damage patterns of terminal deamination common to ancient DNA (Briggs et al., 2007). All our samples present typical damage profiles (Supplementary Figure 1). GEN72, RK1001, Post6 and 1343UnTal85 only retain damage in the last two bases as these libraries were prepared using a ‘UDG-half’ protocol (Rohland et al., 2015, See Methods).

The six reconstructed genomes and their plasmids were compared to the two Bronze Age genomes reported previously (Rasmussen et al., 2015). After visual inspection of aligned reads, our prehistoric genomes from Europe showed similar coverage of the reference genome CO92, and all regions were also covered in the Bronze Age Altai *Y. pestis* genomes (Figure 2A). The six reconstructed genomes in this study lack the same region of the pMT1 plasmid, which contains the *ymt* gene (Figure 2), as already identified in the Altai genomes (Rasmussen et al., 2015). The *ymt* gene codes for the *Yersinia* murine toxin, which is an important virulence factor in *Y. pestis* related to transmission via the flea vector (Hinnebusch et al., 2002, 2000). The expression of *ymt* protects the bacteria from toxic blood digestion by-products in the flea's gut and thus functions to aid in colonization of the flea midgut (Hinnebusch et al., 2002).

**Figure 2:**
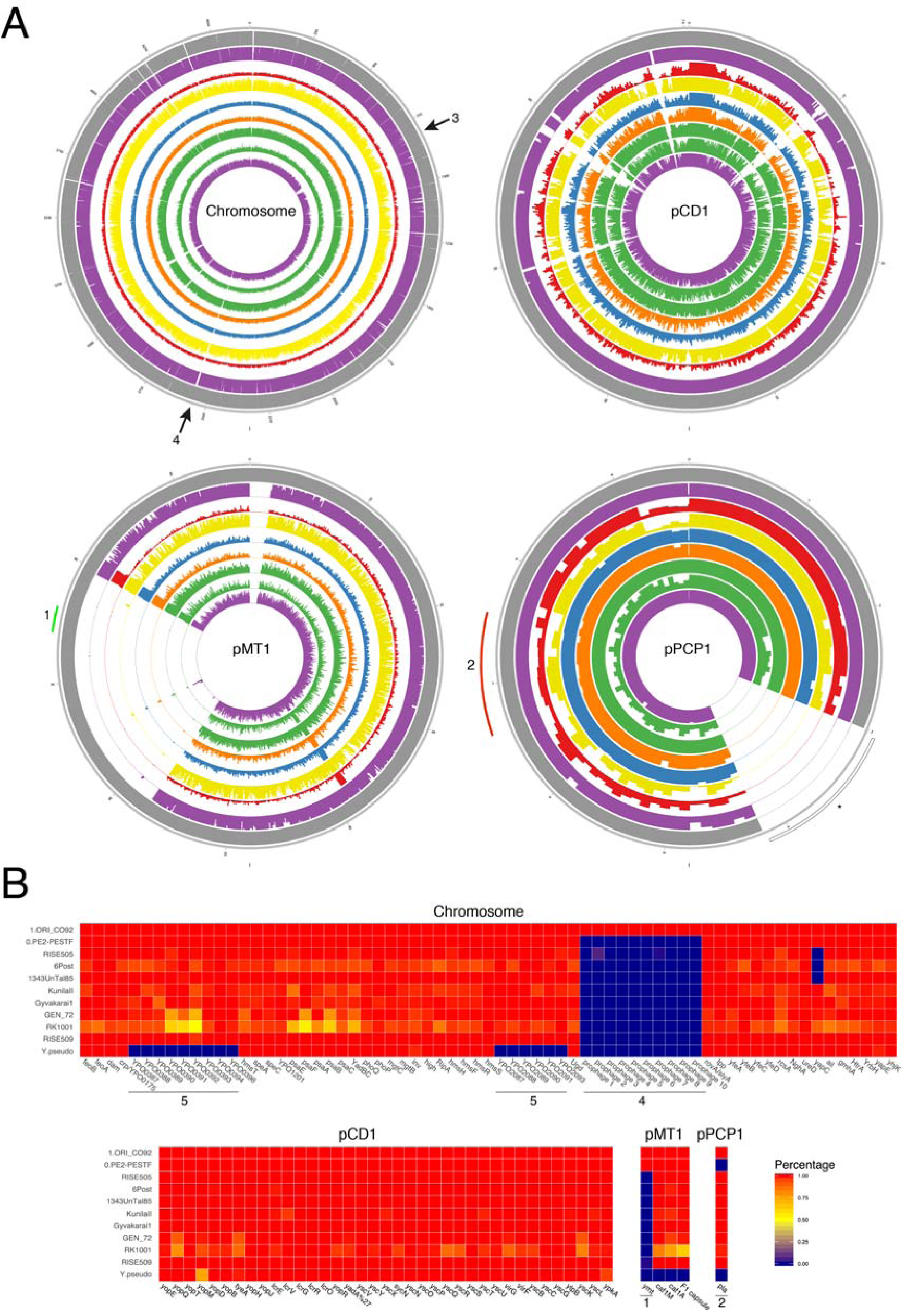
A) Average coverage plot for the chromosome and plasmids of Yersinia pestis, from the outer ring to the inner ring: Y. pestis CO92 (NC_003143.1, reference), RISE509, RK1001, GEN72, Gyvakarai1, KunilaII, 6Post, 1343UnTal85 and RISE505. Colours correspond to the regions where the genomes were recovered from: Altai region (purple), Russia (red), Croatia (dark yellow), Gyvakarai, Lithuania (blue), Kunila, Estonia (orange), Augsburg, Germany (green). The average depth of coverage was calculated for 1kb regions for the chromosome and 100bp for the plasmids, each ring represents a maximum of 20X coverage. The figure was generated with Circos (Krzywinski et al., 2009). B) Percentage covered of virulence factors located on the Yersinia pestis chromosome and plasmids, plotted with in R using the ggplot2 package. [1] ymt gene, [2] pla, [3] deletion of flagelin genes, [4] filamentous prophage YpfΦ, [5] Y. pestis-specific genes, [*] region mask in pPCP1 due to high similarity to expression vectors during enzyme production (Schuenemann et al., 2011).

### Phylogeny and Dating

To assess the phylogenetic positioning of the six European LNBA *Y. pestis* genomes with respect to the modern and ancient *Y. pestis* genomes, Neighbour Joining (NJ, Supplementary Figure 2A), Maximum Parsimony (MP, Supplementary Figure 2B) and Maximum Likelihood (ML, Figure 3, Supplementary Figure 2C) trees were computed. Our samples form a distinct clade in the *Y. pestis* phylogeny together with the previously reconstructed Southern Siberian Bronze Age *Y. pestis* genomes (Rasmussen et al., 2015). This topology has a high bootstrap support of >95% in all three methods. The branching point of the LNBA genomes with the main branch leading towards the modern *Y. pestis* strains represents the most recent common ancestor (MRCA) of all the extant and ancient *Y. pestis* genomes currently available.

**Figure 3:**
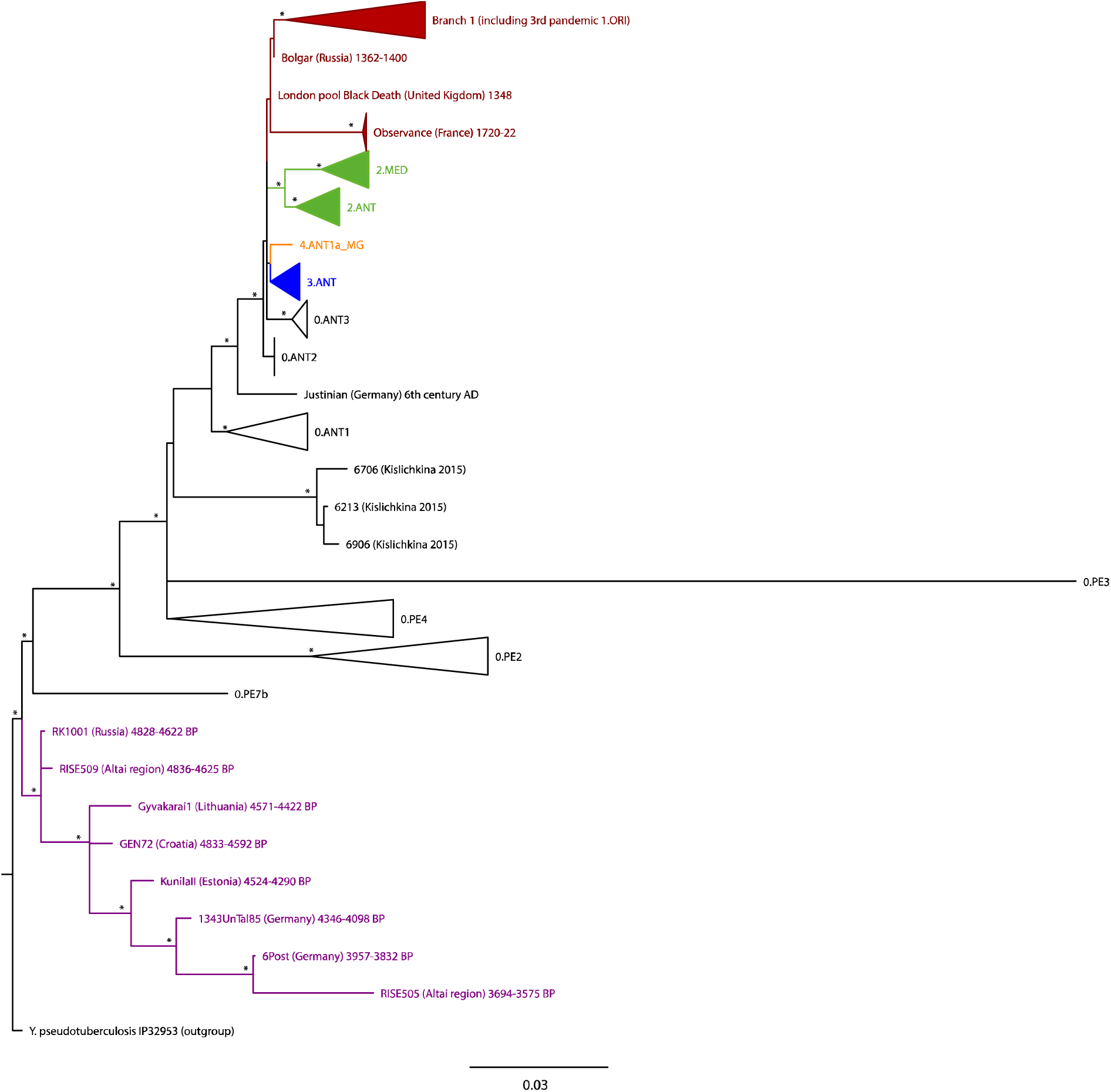
Maximum Likelihood tree of all Yersinia pestis genomes including 1,867 SNPs positions with complete deletion. Nodes with support equal or higher than 95% are marked with an asterisk. The colours represent different branches in the Y. pestis phylogeny: branch 0 (black), branch 1 (red), branch2 (green), branch 3 (blue), branch 4 (orange) and LNBA Y. pestis branch (purple). Y. pseudotuberculosis-specific SNPs were excluded from the tree for representative matters.

To date the MRCA of *Y. pestis* we performed a ‘tip dating’ analysis using BEAST (Drummond et al., 2012). The MRCA of all *Y. pestis* was dated to 6,078 years (95% HPD interval: 5,036-7,494 years) suggesting a Holocene origin for plague, which is in agreement with previous estimates (5,783 years, 95% HPD interval: 5,021–7,022 years, Rasmussen et al., 2015). The time to the MRCA of *Y. pestis* and *Y. pseudotuberculosis* strain IP 32953 was estimated to 28,258 years (95% HPD interval: 13,200-44,631 years). A maximum clade credibility tree was computed (Supplementary Figure 3) supporting the same topology as the NJ, MP and ML with high statistical support of the branching points of the LNBA plague clade.

### Genetic makeup

The effects of Single Nucleotide Polymorphisms (SNPs) detected in our dataset were determined using the software *snpEff* (Cingolani et al., 2012) and an in-house program (*MultiVCFAnalyzer*). A total of 423 SNPs were found in the LNBA branch including strain-specific and shared SNPs. A total of 114 synonymous and 202 non-synonymous SNPs are present in the LNBA branch. All the LNBA genomes share five SNPs: four non-synonymous and one stop mutation (Supplementary Table 2).

Additionally, nine stop mutations were detected in the ancient branch, which were not shared by all the LNBA genomes. Most of these mutations were found in the terminal part of the LNBA branch with six being specific to the youngest Early Bronze Age *Y. pestis* strain (RISE505), one being shared between RISE505 and Post6, one being GEN72-specific and one being Gyvakarai1-specific (Supplementary Table 3). Additionally, RISE505 misses the start codon of the YPO0956 gene, which is involved in iron transport, and one stop codon in YPO2909, which is a pseudogene.

To identify potential homoplasies, a table of all variable SNPs was examined for any that contradict the tree topology. Nine homoplasies were detected (Supplementary Table 4). Furthermore, a tri-allelic site was detected at nucleotide position 4,104,762 (A,T,C).

The percentage of the gene covered in the LNBA plague genomes was calculated for a set of genes that are related to virulence, flea transmission and colonization and dissemination (Figure 2B). We observed the absence of YpfΦ (Derbise et al., 2007), a filamentous prophage, in all LNBA plague genomes. While YpfΦ is found in some *Y. pestis* strains of branch 0, branch 1 and branch 2 as a free phage, it has only been fully integrated and stabilized into the chromosome of the strains 1.ORI which are responsible for the third pandemic (Derbise and Carniel, 2014). Additionally, the *yapC* gene was lost in the three younger LNBA strains (1343UnTal85, Post6 and RISE505). YapC was initially thought to be involved in the adhesion to mammalian cells, autoagglutination and biofilm formation when expressed in *E. coli* (Felek et al., 2008). However, the *yapC* knockout in *Y. pestis* does not affect those functions. Felek and colleagues have thus suggested that this is due to either low expression of *yapC in vitro* or by compensation through other genes (Felek et al., 2008). The only virulence factor located in the plasmids missing in all the LNBA *Y. pestis* strains is *ymt* (Figure 2). Other virulence factors, such as *pla* and *caf1,* were already present in the LNBA*Y. pestis* genomes. The *pla* gene is involved in the dissemination of the bacteria in the mammalian host by promoting the migration of the bacteria to the lymphatic nodes (Lathem et al., 2007; Sebbane et al., 2006), while the *caf1* gene encodes the F1 capsular antigen, which confers phagocytosis resistance to the bacterium (Du et al., 2002). Both genes are absent in the closest relative *Y. pseudotuberculosis*.

Urease D (*ureD*) is an important gene that plays a role in flea transmission. When *ureD* is expressed in the flea vector it causes a toxic oral reaction to the flea killing around 30-40% (Chouikha and Hinnebusch, 2014). While *ureD* is functional in *Y. pseudotuberculosis*, it is a pseudogene in *Y. pestis*. The pseudogenization of this gene is caused by a frameshift mutation (insertion of a G in a six G-stretch) in *Y. pestis* (Sebbane et al., 2001). The LNBA *Y. pestis* genomes were inspected in the search of this specific frameshift mutation. This insertion is not present in those genomes indicating that this gene was still functional in *Y. pestis* at that time, suggesting that it was as toxic to fleas as its ancestor *Y. pseudotuberculosis*.

Large-scale insertions and deletions (indels) were evaluated by comparison of mapped data for the LNBA *Y.* pestis genomes, branch 0 strains (0.PE7, 0.PE2-F. 0.PE3, 0.PE4), KIM, and CO92 using *Y. pseudotuberculosis* IP 32953 (NC_006155.1) as a reference. Regions larger than 1 kb were explored as possible indels. We detected two regions present in the LNBA *Y. pestis* genomes that are absent in all the other strains analyzed: a 1kb region (2,587,386-2,588,553) that contains a single gene (YPTB0714) encoding an aldehyde dehydrogenase, part of the R3 *Y. pseudotuberculosis*-specific region identified by Pouillot et al., 2008 and a second region (1.5kb, 3,295,644-3,297,223) that contains a single gene (YPTB2793) encoding a uracil/xanthine transporter being part of the region orf1 defined by Pouillot et al., 2008, which was also characterized as *Y. pseudotuberculosis*-specific. Additionally, two missing regions were detected: one region of 34kb is missing in the three younger genomes of the LNBA lineage (Post6, 1343UnTal85 and RISE505) and another 36kb region, which contains flagella genes, is missing in the youngest sample RISE505, as shown by Rasmussen et al., 2015, which contains flagella genes. These two missing regions contain multiple membrane proteins, which could be potential virulence factors or antigens recognized by the immune system of the host.

## Discussion

The six prehistoric genomes presented here are the first complete *Y. pestis* genomes spanning from the Late Neolithic to the Bronze Age in Europe. They form a distinct clade with the previously reconstructed Southern Siberian Bronze Age *Y. pestis* genomes, confirming that all LNBA genomes identified so far originate from a common ancestor. The previous reported genome RISE509 (Rasmussen et al., 2015) together with the reconstructed RK1001 genome reported here occupy the most basal position of all *Y. pestis* genomes sequenced to date. This suggests that Central Eurasia rather than Eastern Asia should be considered as the region of potential plague origin.

The temporal and spatial distribution of the Late Neolithic and Bronze Age *Y. pestis* genomes allows us to evaluate the evolution and dissemination of plague in prehistory. We propose two contrasting scenarios to explain the phylogenetic pattern observed in the LNBA *Y. pestis* branch:

1. **Plague was introduced multiple times to Europe** from a common reservoir between 5,000 to 3,000 BP. Here, the bacterium would have been spread independently from a source, most likely located in Central Eurasia, to Europe at least four times during a period of over 1,000 years (Figure 1A), once to Lithuania and Croatia, once to Estonia, and two times to Southern Germany. A similar “multiple wave” proposal has been made for the second pandemic, where climatic fluctuation was considered as driving changes in rodent populations (Schmid et al., 2015). We do not have such data for the time periods in question here, and thus cannot speculate on the mechanism.
2. **Plague entered Europe once during the Neolithic.** From here it established a reservoir within or close to Europe from which it circulated, and then moved back to Central Eurasia and the Altai region/East Asia during the Bronze Age (Figure 1B). This parallels the scenario of local persistence and eastward movement during the second pandemic that is gaining support as more genetic data become available (Seifert et al., 2016; Spyrou et al., 2016).

With just a few genomes available it is difficult to disentangle the two hypotheses; however, interpreting our data in the context of what is known from human genetics and archaeological data can offer some resolution. Ancient human genomic data point to a change in mobility and a large scale expansion of people from the Caspian-Pontic Steppe related to individuals associated with the ‘Yamnaya’ complex, both to the East and the West starting around 4,800 BP. These people carried a distinct genetic component that first appears in Central European individuals from the Corded Ware Complex and then forms/becomes part of the genetic composition of most subsequent and all modern day European populations (Allentoft et al., 2015; Haak et al., 2015). It was furthermore shown that there is a close genetic link between the highly mobile groups of people associated to the Southern Siberian ‘Afanasievo Complex’, the ‘Yamnaya’, and the Central and Eastern European Corded Ware Complex (Allentoft et al., 2015).

Our earliest indication of plague in Europe is found in Croatia and the Baltic region and coincides with the time of the arrival of the genetic steppe component (Allentoft et al., 2015). The two Late Neolithic *Y. pestis* genomes from the Baltic in this study were reconstructed from individuals associated with the Corded Ware Complex (Gyvakarai1 and KunilaII). The Baltic and Croatian *Y. pestis* genomes are genetically derived from a common ancestor of the strain RK1001, reconstructed from an individual associated to the ‘Yamnaya’ complex and RISE509 from the ‘Afanasievo’ complex from the Altai region, suggesting that the pathogen might have spread with steppe people from Central Eurasia to Eastern and Central Europe during their large scale expansion. Furthermore, human genomic analyses indicate that the individuals RISE509, Gyvakarai1, KunilaII and GEN72 carry ‘steppe ancestry’ (Mathieson et al., 2017; Mittnik et al., 2017) Evidence for these long distance contacts is also present in the archaeological record. For example, the Gyvakarai1 burial is characterised by both a specific set of grave inventory (hammer headed pin) and distinct skeletal morphology, which have no analogues in earlier local populations (Tebelškis and Jankauskas, 2006).

The younger Late Neolithic *Y. pestis* genomes from Southern Germany are genetically derived from the Baltic strains and are found in individuals associated with the Bell Beaker Complex. Previous analyses have shown that Bell Beaker individuals from Germany also carry ‘steppe ancestry’ (Allentoft et al., 2015; Haak et al., 2015). This suggests that *Y. pestis* may have been spread further southwestwards analogous to the human steppe component. The youngest of the LNBA *Y. pestis* genomes (RISE505), found also in the Altai region, associated with the Central Eurasian ‘Andronovo’ complex, descends from the Central European strains, which suggests a spread back into Southern Siberia. Interestingly, genome-wide human data shows that human individuals associated to the Sintashta, Srubnaya and Andronovo cultural complexes in the Eurasian steppes (dating from around 3,700-3,500 BP) carried mixed ancestry of middle Neolithic European farmers and Bronze Age steppe people, suggesting a backflow of human genes from Europe to Central Eurasia (Allentoft et al., 2015). Form an archeological perspective there is a close connection of the Abashevo cultural complex and Sintashta, that might also have included population shifts West to East. In particular, the post-Sintashta Andonovo cultural complex is an epoch of massive population shifts affecting all the area east of the Urals to the Western borders of China including populations with European origin (Koryakova and Epimakhov, 2007; Kuzmina, 2008). The steppe, as a natural corridor connecting people and their livestock throughout Central and Western Eurasia, might have facilitated the spread of strains closely related to the European Early Bronze Age *Y. pestis* back to the Altai region, where RISE505 was found. The patterns in human genetic ancestry and admixture, in combination with the temporal series within the LNBA *Y. pestis* branch, therefore support scenario 2, suggesting that *Y. pestis* was introduced to Europe from the steppe around 4,800 BP. Thereafter, the pathogen became established in a local reservoir within or in close proximity to Europe, from where the European *Y. pestis* strain was disseminated back to the Altai region in a process connected to the backflow of human genetic ancestry from Western Eurasia into Southern Siberia. The pathogen diversity, therefore, mirrors the archaeological evidence, which indicates a strong intensification of Eurasian networks since the beginning of the Bronze Age (Vandkilde, 2016).

Even though *Y. pestis* seems to have been spread following human movements, its mode of transmission during this early phase of its evolution cannot be easily determined. Most contemporary cases of *Y. pestis* infection occur via an arthropod vector and stem from a sylvatic rodent population that has resistance to the bacterium. The flea transmission can be accomplished by one of two mechanisms: the classical blockage-dependent flea transmission (Hinnebusch et al., 1998) and the recently proposed early-phase transmission (EPT) (Eisen et al., 2006). In the blockage-dependent model, *Y. pestis* causes an obstruction in the flea digestive system by producing a biofilm that blocks the pre-gut of the flea within 1-2 weeks after infection. This blockage prevents a blood meal from reaching the flea's gut, and regurgitation of the blood by a hungry flea in repeated attempts to feed sheds several live bacteria into the blood stream of the host (Chouikha and Hinnebusch, 2012; Hinnebusch et al., 1998). It has been shown that the blockage-dependent transmission requires a functional *ymt* gene and *hms* locus, and non-functional *rcsA, pde2* and *pde3* genes (Sun et al., 2014). *ymt* protects *Y. pestis* from toxic by-products of blood digestion and allows the bacterium to colonise the mid-gut of the flea. The *hms* locus is involved in biofilm formation and *rcsA*, *pde2* and *pde3* are the down-regulators of biofilm formation. However, evidence is emerging that *Y. pestis* can be transmitted efficiently within the first 1-4 days after entering the flea prior to biofilm formation (Eisen et al., 2015, 2006), in a process known as the EPT model. Unfortunately this model is currently less well understood molecularly and physiologically than blockage-dependent transmission, but has been shown to be biofilm (Vetter et al., 2010) and *ymt* independent (Johnson et al., 2014).

Based on the genetic characteristics of the LNBA genomes (i.e. lack of *ymt*, still functional *pde2* and *rcsA* as shown by previous work (Rasmussen et al., 2015), functional *ureD* which will kill 30-40% of the flea vectors) it seems most parsimonious that *Y. pestis* was not able to use a flea vector in a blockage-dependent model. However, since none of these genes seem to be required for EPT, it remains possible that LNBA *Y. pestis* was transmitted by a flea vector via this transmission mode. Under this assumption, the transmission would have been presumably less efficient since a functional Urease D would have reduced the number of fleas transmitting the bacteria.

The presence of genes involved in virulence in the mammalian host such as *pla* and *caf1*, which are absent in *Y. pseudotuberculosis*, indicates that LNBA *Y. pestis* was already adapted to mammalian hosts to some extent. *pla* aids in *Y. pestis* infiltration of the mammalian host (Lathem et al., 2007; Sebbane et al., 2006). The *pla* gene present in the LNBA *Y. pestis* strains has the ancestral I259 variant, which has been shown to be less efficient than the derived T259 form (Haiko et al., 2009). *Y. pestis* with the ancestral variant is able to cause pneumonic disease, however, it is less efficient in colonising other tissues (Zimbler et al., 2015). This indicates that LNBA *Y. pestis* could potentially cause a pneumonic or a less virulent bubonic form. In addition to the above noted changes, we detected two regions missing in the LNBA genomes: a ∼34kb region that contains genes encoding membrane proteins missing in the three youngest *Y. pestis* strains (Post6, 1343UnTal85 and RISE505) and a ∼36kb region containing genes encoding proteins involved in flagellin production and iron transporters missing in the youngest sample RISE505, as observed elsewhere (Rasmussen et al., 2015). This genome decay affecting membrane and flagellar proteins potentially involved in interactions with the host's immune system, can be an indication of adaptation to a new host pathogenic lifestyle (Ochman and Moran, 2001).

Our common understanding is that plague is a disease adapted to rodents, where commensal species such as *Rattus rattus* and their fleas play a central role as disease vectors for humans (Perry and Fetherston, 1997). While a rodent-flea mediated transmission model is compatible with the genomic makeup of the LNBA strains, disease dynamics may well have differed in the past. The most parsimonious explanation would be that LNBA plague indeed traveled with rodent species commensal to humans, in keeping with the orthodox model of plague transmission. The Neolithic is conventionally considered to be a time period where new diseases were introduced into human groups as they made the transition from a nomadic lifestyle to one of sedentism, and where the adoption of agriculture and increased population density acted synergistically to change the disease landscape (Ronald Barrett et al., 1998). Whether commensal rodent populations were large enough to function as reservoir populations for plague during human migrations at this time is unknown. In central Eurasian Bronze Age cultures, agriculture, i.e. large scale food storage, is mostly absent (Ryabogina and Ivanov, 2011) However, contact between steppe inhabiting rodents, pastoralists and their herds might have been frequent when moving within these environments. Alternative models of transmission involving different host species, perhaps even humans or their domesticates, might carry some traction, as the ancient disease may have behaved rather differently from the form we know today.

Here, we present the first LNBA *Y. pestis* genomes from Europe. We show that all LNBA genomes reconstructed so far form a distinct lineage that potentially entered Europe following the migration of steppe people around 4,800 BP. We find striking parallels between the *Y. pestis* dispersal pattern and human population movements during this time period. We propose two scenarios for presence of the bacteria in Europe: a multiple introduction hypothesis from a Central Eurasian source, or the establishment of a local *Y. pestis* focus within or close to Europe from where a resident strain ultimately moved back towards Central Asia in the Bronze Age. On account of the chronology and the tight synergy between the ancient *Y. pestis* phylogeny and known patterns of human mobility, we find stronger support for the second scenario.

The LNBA period was a time of increased mobility and cultural change. The presence of *Y. pestis* may have been a promoting factor for the increase in mobility of human populations (Rasmussen et al., 2015). The manifestation of the disease in Europe could have played a major role in the processes that led to the genetic turnover observed in the European human populations, who may have harbored different levels of immunity against this newly introduced disease. Testing these hypotheses will require more extensive assessment of both human and *Y. pestis* genomes from the presumed source population before and after migration from the steppes, as well as in Europe during this period of genetic turnover.

## Authors contribution

J.K., A.H. and A.A.V. conceived the study. K.M., R.A., M.D., R.J., M.T., P.W.S., A.B., I.J., M. N., S.R., M.S., A.S., S.H. provided the samples and performed archaeological assessment. A.M., S.P., M.F., A.A.V. performed laboratory work. A.A.V., A.H., M.A.S., F.M.K. and J.K. analysed the data. A.A.V, A.H., J.K., P.W.S., K.I.B., W.H. and A.M. wrote the manuscript with contributions from all co-authors. All authors read and approved the final manuscript.

### Data availability

Raw sequencing data have been deposited at the European Nucleotide Archive under accession PRJEBXXXXX

## Acknowledgements

We thank Corina Knipper, Ernst Pernicka, Stephanie Metz, Fabian Wittenborn, Stephan Schiffels, Joris Peters, Michaela Harbeck, and all the members of the Archaeogenetics Department of the Max Planck Institute for the Science of Human History for helpful discussion and suggestions. We thank Annette Günzel for graphical support. We thank Isil Kucukkalipci, Antje Wissgott, Marta Burri and Franziska Göhringer for technical support in the lab. We thank Prof. Dr. Joachim Wahl and Dr. Gunita Zatina and Prof. Andrejs Vasks for kindly providing the Althausen samples and the Latvian samples, respectively, used in this study. We also thank Josip Burmaz and Dženi Los for providing the Croatian sample. We thank Natalia Berezina and Dr. Julia Gresky for providing an anthropological assessment of RK1001. We thank James A. Fellows Yates for proof-reading the manuscript.

The genetic and archaeological research on the human individuals from the Augsburg region was financed by the Heidelberg Academy of Science within the WIN project “Times of Upheaval: Changes of Society and Landscape at the Beginning of the Bronze Age”. This work was also supported by the Max Planck Society and the European Research Council starting grant APGREID (to J.K.). M.N. and I.J. were supported by the Croatian Science Foundation grant [1450].

## STAR Methods

### Sampling and extraction

Sampling of a total of 563 tooth and bone samples (Russia (122), Hungary and Croatia (139), Lithuania (27), Estonia (45), Latvia (10), and Germany (Althausen 4, Augsburg 83, Mittelelbe-Saale 133)) took place in the clean room facilities of the Institute for Archaeological Sciences at the University of Tu□bingen, the Institute of Archaeology RCH HAS in Budapest and of the MPI-SHH in Jena. After irradiation with UV light to remove surface contamination, teeth were sawed apart transversally at the border of crown and root, and dentine from inside the crown was sampled and powdered using a sterile dentistry drill. For the samples processed in Budapest, whole teeth were powdered. For bone samples, the surface layer from the sampling area was removed with a dentistry drill prior to obtaining bone powder from the inside of the bone by drilling. For each specimen we gathered between ∼30 and 120 mg of powder to be used for DNA extraction.

Extraction was performed following a protocol optimized for the recovery of small ancient DNA molecules (Dabney et al., 2013), resulting in 100μl of DNA extract per sample. An aliquot of 20μl of extract was used to generate double-indexed libraries (Kircher et al., 2012; Meyer and Kircher, 2010). Negative controls were included in the extraction and library preparation and taken along for all further processing steps.

### Shotgun screening

Libraries were PCR-amplified and quantified using an Agilent 2100 Bioanalyzer DNA 1000 chip and pooled at equimolar concentrations prior to paired-end sequencing on a NextSeq500 with 2×101+8+8 and a HiSeq2500 with 2×101+8+8 cycles according to the manufacturer's instructions to a depth of ∼1.5 million reads per library.

### *In-silico* screening

The sequencing data for the 170 samples was preprocessed with ClipAndMerge (Peltzer et al., 2016) to remove adaptors, base quality-trim (20) and merging and filtering for only merged reads. Reads were mapped using the BWA aln algorithm (Li and Durbin, 2009) to a multi-species reference panel, containing various representatives of the genus *Yersinia* (Table 2) and the plasmids of *Yersinia pestis*: pCD1, pMT1 and pPCP1 from *Y. pestis* CO92. The region comprising 3000-4200bp of the Y. pestis specific plasmid pPCP1 was masked in the reference, since it is highly similar to an expression vector used during the production of enzyme reagents (Schuenemann et al., 2011).

Mapped files were then filtered for reads with a mapping quality higher than 20 with Samtools (Li et al., 2009). PCR duplicates were removed using the MarkDuplicates tool in Picard (1.140, http://broadinstitute.github.io/picard/). The number of reads mapping specifically to each genome and to the plasmids were retrieved from the bam files using Samtools (Li et al., 2009) idxstats. An endogenous based score was used to assess the potential of the sample being ‘positive’ for *Y. pestis*. It was calculated as follows:

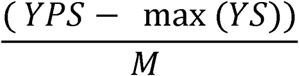

where YPS is the number of reads specifically mapping to *Y. pestis*; YS is the maximum number of reads mapping specifically to a *Yersinia* species with the exception of *Y. pestis* and M is the total number of merged reads in the sample. By using the maximum number of reads mapping to another species of the genus *Yersinia*, the score takes in account different source of contamination other than *Y. pseudotuberculosis*. Five samples (RK1001, Gyvakarai1, Kunilall, 6Post and 1343UnTal85) fulfilled the criteria for being considered strong candidates (score higher than 0.005 and reads mapping to all plasmids). Another samples, GEN72, was also included in further processing and analysis since it had higher numbers mapping the *Y. pestis* chromosome and plasmids even though it did not full-fill the score requirements. For a detailed description of the archaeological sites and individuals see the SI.

### Deep shotgun sequencing

The five strong candidate samples detected in screening of the shotgun data were processed for deep shotgun sequencing as following: For Gyvakarai1 the screening library described above was pair-end sequenced on two lanes of a HiSeq4000 for 100 cycles, and on a full run of a NextSeq500 for 75 cycles. The screening library for KunilaII was pair-end sequenced deeper on 80% of one lane of a HiSeq4000 for 100 cycles. Additionally, 40 μl of DNA extract of KunilaII was converted in to a library treated with UDG and endonuclease VIII to remove deaminated bases (Briggs and Heyn, 2012), and pair-end sequenced on one lane of a HiSeq4000 for 75 cycles.

For RK1001, Post6 and 1343UnTal85, 60 μl of DNA extract each were converted into DNA libraries using so-called UDG-half treatment, whereby deaminated bases are partially removed and retained mostly at the ends of the molecule (Rohland et al., 2015). The library of RK1001 was deep shotgun pair-end sequenced in 8 lanes of a HiSeq4000 for 55 cycles. The libraries of 6Post and 1343UnTal85 were deep shotgun single-end sequenced on 2 and a half lanes of a HiSeq4000 for 75 cycles. Post6 was additionally pair-end sequenced on a full run of a NextSeq500 for 75 cycles.

### *Y.pestis* in-solution capture

*Y. pestis* whole-genome DNA capture probes were designed using as template sequences the *Y. pestis* CO92 chromosome (NC_003143.1), *Y. pestis* CO92 plasmid pMT1 (NC_003134.1), *Y. pestis* CO92 plasmid pCD1 (NC_003131.1), *Y. pestis* KIM 10 chromosome (NC_004088.1), *Y. pestis* Pestoides F chromosome (NC_009381.1) and Y. pseudotuberculosis IP 32953 chromosome (NC_006155.1). We used a 6 bp tiling with a probe length of 52 bp with an additional 8 bp 3′ linker sequence as described in (Fu et al., 2013). Low complexity regions were masked using dustmasker (Camacho et al., 2009, version 2.2.32+). Redundant probes as well as probes with more than 20% masked nucleotides were discarded. The procedure resulted in 816,413 unique probe sequences. A second probe set was created with a coordinate offset of 3 bp resulting in 827,438 unique probe sequences. In combination the two probe sets represent an effective tiling density of 3 bp. The two probe sets were ordered on two 1 million feature Agilent SureSelect DNA Capture Arrays. The full capacity of the arrays was filled up with randomly selected probes. The two arrays were turned into in-solution DNA capture libraries as described elsewhere (Fu et al., 2013).

For GEN72, 25 μl of DNA extract was converted into DNA libraries using so-called UDG-half treatment as described above^10^. The UDG-half libraries of RK1001 and GEN72 were enriched for *Y. pestis* DNA using in-solution DNA capture probes (see above) as described elsewhere (Fu et al., 2013; Haak et al., 2015; Mathieson et al., 2015). The capture products of RK1001 and GEN72, were sequenced on 1 and 0.6 of the lane, respectevely, of the HiSeq4000 for 75 cycles.

### Genome reconstruction and authentication

All samples were processed with the EAGER pipeline (Peltzer et al., 2016). Sequencing quality for each sample was evaluated with FastQC (http://www.bioinformatics.babraham.ac.uk/projects/fastqc/), and adaptors clipped using the ClipAndMerge module in EAGER. For paired-end data, the reads were also merged with ClipAndMerge and only the merged reads were kept for further analysis.

Due to variability in the laboratory preparation and sequencing strategies, the sequencing reads for each sample were treated as follows:

- Gyvakarai1: two HiSeq lanes and one Next-Seq run paired-end of the non-UDG treated library were combined and reads mapped to *Y. pestis* CO92 reference with BWA aln (−l 16, −n 0.01, hereby referred to as non-UDG parameters). Reads with mapping quality scores lower than 37 were filtered out. PCR duplicates were removed with MarkDuplicates. MapDamage (Jónsson et al., 2013, v2.0) was used to calculate damage plots. Coverage was calculated with Qualimap (Okonechnikov et al., 2016, v2.2).
- KunilaII: UDG and the non-UDG libraries were sequenced in 2 HiSeq pair-end lanes and processed separately until calculation of the coverage. The non-UDG treated libraries were mapped with non-UDG parameters while the UDG treated library reads were mapped with more stringent parameters (-l 32, -n 0.1, referred to as UDG parameters). Reads with mapping qualities less than 37 were filtered out and duplicates were removed with MarkDuplicates as before. The non-UDG bam file was used to calculate damage plots as indicated above. After duplicate removal, the UDG- and non-UDG treated BAM files were merged together and used to calculate the coverage as above.
- GEN72, Post6 and 1343UnTal85: the UDG-half treated libraries were sequenced in two HiSeq lanes for Post6 and 1343UnTal85 and 19,777,683 reads were generated in the HiSeq for GEN72, and two different runs were performed. For the first run, reads without clipping were used to retain miscoding lesions indicative of aDNA. BWA aln was used for mapping with non-UDG parameters (−l 16 and −n 0.01). Reads with mapping qualities lower than 37 were filtered and PCR duplicates were removed with MarkDuplicates as described above. Coverage and damage plots were calculated as above. After clipping the last two bases with the module ClipAndMerge in eager, potentially affected by damage, the samples were mapped with UDG parameters.
- RK1001: UDG-half library was shotgun sequenced pair-end in 8 HiSeq lanes and in-solution captured and sequenced single end to a depth of 303,148,884 reads sequenced in the HiSeq. Shotgun and captured data were combined in a fastq file and processed as described above for GEN72, Post6 and 1343UnTal85.

### SNP calling & phylogenetic analysis

Prior to SNP calling in order to avoid false SNP calling due to aDNA damage, the quality scores of damaged sites in the non-UDG treated samples were downscaled using MapDamage (Jónsson et al., 2013, v2.0), as performed in previous analysis (Rasmussen et al., 2015). For the UDG-half data, the files with the two last bases clipped, hence removing the damage signal, and mapped with UDG parameters were used for SNP calling (see above). SNP calling was performed with GATK UnifiedGenotyper (Van der Auwera et al., 2013) in EAGER^43^ with default parameters and the ‘EMIT_ALL_SITES’ output mode.

VCF files of the new ancient samples, along with the two complete genomes from Rasmussen et al., 2015, the Black Death (Bos et al., 2011), Justinianic Plague (Feldman et al., 2016), Bolgar (Spyrou et al., 2016) and Observance (Bos et al., 2016) genomes, were combined with a curated dataset of 130 modern genomes (Cui et al., 2013) in addition to 11 samples from the Former Soviet Union (Zhgenti et al., 2015) and 19 draft genomes of *Y. pestis* subsp. *microtus* strains (Kislichkina et al., 2015).

The VCF files were processed with an in-house program (*MultiVCFAnalyser*) that produced a SNP table and an alignment file containing all variable positions in the dataset, in respect to the reference *Y. pestis* CO92. In order to call a SNP a minimum genotyping quality (GATK) of 30 was required, with a minimum coverage of 3X, and with a minimal allele frequency of 90% for a homozygous call. No heterozygous calls were included in the output files.

The SNP alignment was curated by removing all alignment columns with missing data (complete deletion). The curated SNP alignment was then used to compute NJ and MP trees with MEGA6 (Tamura et al., 2013) and a ML tree using PhyML 3.0 (Guindon et al., 2010) with the GTR model used in previous *Y. pestis* work (Cui et al., 2013; Rasmussen et al., 2015), with 4 gamma categories and the best of NNI and SPR as tree branch optimization. The specific variants of *Y. pseudotuberculosis* were removed from the analysis to improve the visual resolution of the tree.

### Dating analysis

The SNP alignment after complete deletion was used for molecular dating using BEAST 1.8.2 (Drummond et al., 2012). The modern sample 0.PE3, also called Angola, was removed from the dataset due to its long branch.

For tip dating, all modern genomes were set to an age of 0. The dates of the ancient samples presented in this study plus the two complete genomes from Rasmussen et al., 2015 were recalibrated with Calib 7.1 (http://calib.qub.ac.uk/calib/) to the IntCal13 calibration curve. The ancient samples were given the median calibrated probability as their age, and the 2 sigma interval was used as the boundaries for a uniform prior sampling (Supplementary Table 5). The dates published for previous historical genomes were transformed to cal BP assuming 1950 as age 0 and given the mean as the age with the interval as the boundaries of a prior uniform distribution: Black Death 603 (602-604, Bos et al., 2011); Observance 229 (228-230, Bos et al., 2016), Bolgar 569 (550-588, Spyrou et al., 2016) and Justinian 1453 (1382-1524, Feldman et al., 2016).

The molecular clock was tested and rejected using MEGA6. Therefore, we followed previous work and used an uncorrelated relaxed clock with lognormal distribution (Cui et al., 2013; Rasmussen et al., 2015) with the substitution model GTR+G4. Tree model was set up to coalesce assuming a constant population size and a rooted ML tree was provided as a starting tree. Two independent 1,000,000,000 MCMC chains were computed sampling every 5,000 steps. The two chains were then combined using LogCombiner from BEAST 1.8.2 (Drummond et al., 2012) with a 10 percent burn-in (100,000,000 steps per chain). The ESS of the posterior, prior, treeModel.rootHeight, tMRCA_allpestis are 4,589, 4,087, 1,054 and 7,571 respectively. The trees files for the 2 chains were combined with LogCombiner with 100,000,000 of burning and resampled every 20,000 steps giving a total number of 90,000 trees, that were used to produce a Maximum Clade Credibility tree using TreeAnnotator from BEAST 1.8.2 (Drummond et al., 2012).

### SNP effect analysis and virulence factors analysis

The SNP table from *MultiVCFAnalyzer* was provided to *SnpEff* (Cingolani et al., 2012) and the effect of the SNPs within genes present in the dataset was evaluated. Additionally the SNP table was manually assessed for possible homoplasies.

For the virulence factors, the samples were mapped as indicated above but without applying quality filtering and the percentage of coverage was calculated for each region using bedtools (Quinlan and Hall, 2010) and plotted using the package ggplot2 (Wickham, 2009) in R (R Development Core Team, 2008). Additionally, *ureD* was manually explored for SNPs using IGV (Thorvaldsdóttir et al., 2013).

### Indel analysis

The samples including the two complete Bronze Age genomes (Rasmussen et al., 2015) were mapped against *Y. pseudotuberculosis* IP 32953 with bwa with non-UDG parameters (−n 0.01, −l 16), except for RK1001, GEN72, 1343UnTal85 and 6Post that were mapped with bwa with UDG parameters (−n 0.1, −l 32),. The modern genomes from branch 0 (0.PE7, 0.PE2, 0.PE3 and 0.PE4), *Y. pestis* CO92 and *Y. pestis* KIM10 were *in-silico* cut in 100 bp fragments with 1bp tiling and mapped to *Y. pseudotuberculosis* reference using bwa with UDG parameters (−n 0.1, −l 32). The non-covered regions were extracted using the bedtools genomecov function. Missing regions larger than 1kb were comparatively explored in order to identify indels. Using the bedtools intersect function, we extracted regions missing in the Neolithic genomes and present in the modern ones and also the regions missing in the modern ones but still present in the Neolithic genomes. The results were check by manual inspection in IGV (Thorvaldsdóttir et al., 2013).

